# The Spatial Mixing of Genomes in Secondary Contact Zones

**DOI:** 10.1101/016337

**Authors:** Alisa Sedghifar, Yaniv Brandvain, Peter Ralph, Graham Coop

## Abstract

Recent genomic studies have highlighted the important role of admixture in shaping genome-wide patterns of diversity. Past admixture leaves a population genomic signature of linkage disequilibrium (LD), reflecting the mixing of parental chromosomes by segregation and recombination. The extent of this LD can be used to infer the timing of admixture. However, the results of inference can depend strongly on the assumed demographic model. Here, we introduce a theoretical framework for modeling patterns of LD in a geographic contact zone where two differentiated populations are diffusing back together. We derive expressions for the expected LD and admixture tract lengths across geographic space as a function of the age of the contact zone and the dispersal distance of individuals. We develop an approach to infer age of contact zones using population genomic data from multiple spatially sampled populations by fitting our model to the decay of LD with recombination distance. We use our approach to explore the fit of a geographic contact zone model to three human population genomic datasets from populations along the Indonesian archipelago, populations in Central Asia and populations in India.

## 1 Introduction

Populations frequently undergo periods of relative isolation that are followed by secondary contact. During isolation, the evolutionary processes of genetic drift, mutation, and selection act to differentiate populations at many markers throughout the genome. When these populations come back into contact, the restoration of gene flow generates admixed populations, which start as an assemblage of differentiated parental genomes that are broken up every generation by segregation and recombination between chromosomes.

Under this process, linked alleles of the same ancestry will tend to be co-inherited until separated by recombination. Because the parental populations are differentiated with respect to each other, this co-inheritance leads to a nonrandom association of alleles, referred to as linkage disequilibrium (LD). This admixture-induced LD (or admixture-LD) initially extends over a much larger genomic scale than LD does in either parental population and is a signature of relatively recent admixture (Chakraborty and Weiss 1988; Cavalli-Sforza and Bodmer 1971). One can also think of this signature as the persistence of parental haplotypes in admixed populations which, rather than being measured directly, is measured as the extent of co-occurrence along a chromosome of alleles that are diagnostic of parental origin. Recombination acts every generation to gradually break apart long tracts of ancestry into smaller tracts, and so the association between nearby alleles lasts many generations. The physical scale over which admixture-LD breaks down is determined by the timescale over which parental populations have been interbreeding; the conservation of many ancestral haplotypes over large physical distances would imply very recent admixture, whereas a longer history of admixture produces many smaller parental tracts.

Data from many (potentially weakly) differentiated markers allows for the identification and quantification of admixture in individuals (e.g. Pritchard *et al*. 2000) and the inference of the ancestral origin of a given chromosomal region (e.g. Falush *et al*. 2003; Price *et al*. 2009; Hellenthal *et al*. 2014). The continued mixing of differentiated genotypes, as described above, produces predictable population genomic patterns that change through time, and these signals can be used to not only detect past admixture in extant population, but also to learn about the timing and history of these admixture events (e.g. Hellenthal *et al*. 2014; Loh *et al*. 2013; Harris and Nielsen 2013). Such inferences have been used to reconstruct historical population movements, highlighting the importance of admixture in shaping patterns of diversity in human populations (Hellenthal *et al*. 2014; Reich *et al*. 2009; Patterson *et al*. 2012; Loh *et al*. 2013; Moorjani *et al*. 2013). These studies have utilized powerful methods that first identify stretches of chromosome inherited from a particular parental population (admixture tracts Gravel 2012; Hellenthal *et al*. 2014), or measure the covariance, over spatial scales, of variants that are diagnostic of parental populations (admixture-LD Patterson *et al*. 2012; Loh *et al*. 2013), and then infer the genetic scale over which this measured coancestry decays. Commonly this is done by assuming a model of admixture in which one isolated population is formed by a single admixture event in time, with subsequent random mating. Under this simple model, the distribution of admixture tract lengths and the decay of admixture-LD with respect to genetic distance is approximately exponential, with the rate parameter corresponding to the time in generations since admixture. However, violations of the assumptions of the single-pulse model can result in substantial departure between expected and observed rates of decay of coancestry with respect to time.

Models incorporating multiple admixture times, or sustained, migration (Pool and Nielsen 2009; Gravel 2012; Liang and Nielsen 2014; Hellenthal *et al*. 2014) have been built to address more complex admixture scenarios in single populations. However, these do not incorporate the fact that admixture often occurs in a geographic context – beginning at a given point in time, then spreading across space. Most current models treat each admixed population as an independent event, not accounting for this spatial context, even when admixture in spatially distributed populations are potentially attributable to a single historical event.

In this paper we build an alternative model of diffusion of ancestry across geography in time. Specifically, we consider a scenario in which two populations spread back into contact, generating a gradient of admixture across space with the greatest degree of admixture at the point of initial contact. We refer to this mixture across space, where migration is sustained through both time and space, as a contact zone. This geographic mixing leads to departures from a simple model of exponential decay of admixture-LD as there is exchange of migrants between neighboring populations with different admixture proportions. We describe the expected ancestry-LD in contact zones accounting for migration in continuous space. This model provides a framework to simultaneously examine admixture patterns over a set of geographically distributed populations, and a potential geographic null model for studying historical movements of populations. Inference under this model provides a means to estimate both the time at which populations spread back into contact, as well as some measures of dispersal. We analyze several potential human contact zones under our model and show that simpler ‘point’ models of admixture can infer unreasonably recent admixture dates.

## 2 Methods

### 2.1 Outline of neutral model

Consider two differentiated populations along a transect in space, formerly separated by a barrier that completely prevented migration (at position *x* = 0) that was removed *τ* generations ago (Fig. 1). We imagine the barrier as a physical obstruction to migration; however, in practice the two previously isolated populations could come into contact through a variety of means. We use a continuous-space limit of randomly mating (Wright-Fisher) populations on a line, made formal in e.g. Shiga (1980) that can be described informally as follows:

**Figure 1:**
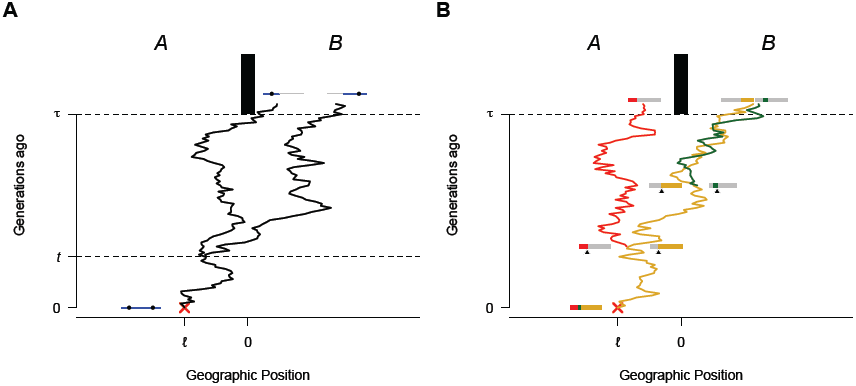
**A:** We follow backward in time the Brownian motion paths of two initially linked lineages, represented here by two black circles located on a grey chromosome. The paths of the two lineages are identical until the first recombination event between them at time *t*, after which they follow independent Brownian paths. The red cross indicates the position, relative to the center of the zone, where the chromosome was sampled in the present day. The black rectangle represents a barrier to dispersal that was removed at time *τ*. In this example, both alleles are of ancestry *B*, since they are on the same side of the barrier to dispersal at time *τ*. **B:** Brownian motion paths of a tract of chromosome. As in Fig. 1**A**, the path along chromosomal fragments are identical until recombination breaks the fragments up. Here, the position of each chromosomal fragment at time *τ* is shown. For the entire portion of chromosome to be of uniform ancestry, all products of recombination must be on the same side of the barrier to dispersal at time *τ*. Here, the green and yellow fragments constitute an unbroken tract of B ancestry.

Since time *τ*, individuals have moved without restrictions following a Gaussian dispersal kernel, in such a way that the distribution of displacements between an ancestor and descendant separated by *t* generations is Gaussian with mean zero and variance *σ*^2^*t*. This forms a gradient of admixed populations across space, whose degree of admixture depends on the time that has passed and the distance to the point of initial contact. Over time, genotypes of different ancestries diffuse across the entire range, and recombination breaks down tracts of continuous ancestry. We aim to describe this diffusion of ancestry throughout time and space.

To determine the typical degree of admixture at a location, we follow the lineage of a sampled individual back through time, tracing the spatial location of the ancestor of today’s sample back to the initiation of secondary contact. The ancestral type of today’s sample is determined by the geographic position of its ancestor *τ* generations ago: we say that a sampled individual whose lineage falls to the left of the barrier (i.e. some point where *x <* 0) is of ancestry *A*, and is otherwise of ancestry *B*. This represents the alleles belonging to ancestral population *A* or *B* before the initiation of secondary contact. We treat time and space as continuous variables, and the time-reversible properties of Brownian motion allow us to model the movement of lineages as a continuous Brownian process.

### 2.2 Behavior of a single locus

We start by describing the properties of a single lineage, *𝒜*, that is sampled at position *ℓ* relative to the center of the contact zone (at *x* = 0), *τ* generations after initial contact. Since we assume the movement of the lineage to be Brownian, the probability that *𝒜* is of ancestry *B* is equal to the probability that the Brownian motion begun at *x* is to the right of zero after *τ* generations, i.e.

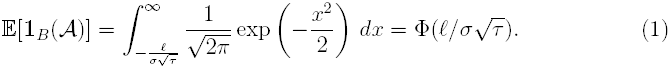

Here 1_*B*_(*𝒜*) is the indicator function:

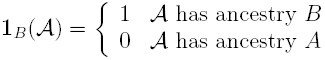

Eq. 1, follows from the assumption that the displacement between parents and off-spring is Gaussian with variance *σ*^2^, allowing us to describe the movement of the lineage after *τ* generations by the Brownian process *B*_*τ*_. The probability then that an individual sampled at geographic position *ℓ* inherits at a given locus from ancestral population *B* is the probability that *x*_*τ*_ > 0 where *x*_*τ*_ ~ *𝒩*(ℓ, *τσ*^2^). This is also the expected frequency of ancestry *B* at position *ℓ*, *τ* generations after contact, and provides an expectation of the cline in ancestry proportion. Although this derivation assumes continuous time, the expression also holds in the case of non-overlapping generations since, if dispersal is Gaussian, the position of an allele at time *τ* is similarly described by a normal distribution.

Under this model, we expect the zone of significant admixture to extend over distance roughly 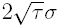 in either direction so, to fit our model using the inference framework we describe below, we will need samples on this spatial scale.

### 2.3 Ancestry LD between linked loci

In our model, all chromosomes begin as unbroken tracts of ancestry prior to initial contact. As time progresses, recombination between haplotypes of different ancestry breaks down these associations. To model this effect, we consider two linked loci separated by a recombination fraction *r*, on a single chromosome sampled at geographic position *ℓ* (see Fig. 1 and legend), and denote the ancestral lineages at these to loci as *𝒜*_1_ and *𝒜*_2_, respectively. The recombination fraction between the loci is the per generation probability of observing a recombinant haplotype as the product of meiosis. For close pairs of markers it may suffice to use the genetic distance *d* in Morgans that separates markers, but for more distant markers we use the probability of an *observed* recombination event, which is the probability of an odd number of recombination events between focal loci, accounting for interference when possible.

We measure ancestry-LD as the covariance in ancestry between the alleles at the two loci

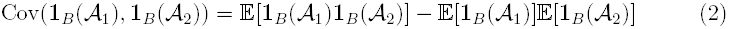

Since *𝒜*_1_ and *𝒜*_2_ are exchangeable, the second term is simply 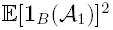, which by Eq. 1 is 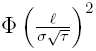.

The first term of Eq. 2 is the probability that both *𝒜*_1_ and *𝒜*_2_ are of ancestry B, which we can compute by considering the Brownian movement of the two lineages. At the time of sampling, and until the first recombination event between the two loci, the two lineages follow an identical path back through time. We assume that after the first recombination event the two lineages never coalesce back onto the same chromosome and therefore pursue independent Brownian paths for the remaining time back to *τ* (Fig. 1). This assumption ignores drift since secondary contact.

This assumption of no drift will be good if 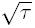 is much smaller than Wright’s neighborhood size *N*_*σ*_, i.e. the number of individuals within a region of width *σ* (Wright 1943). This is because in one dimension, assuming Gaussian dispersal, the number of generations that two randomly moving lineages that start in the same place spend within distance *σ* of each other across *τ* generations is of order 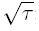; the chance that they coalesce each time they are is proportional to 1*/N*_*σ*_, and so the chance of coalescence is negligible if 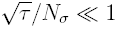. (For more discussion of scaling see e.g. Barton *et al*. (2002).)

To find an expression for this covariance, observe that the random time *T* since the first recombination event between the two loci is exponentially distributed with rate parameter *r*. Given that the most recent recombination along this lineage occured *T* generations ago, with *T* < *τ*, the joint spatial positions (*X*_1_,*X*_2_) of the two lineages (*𝒜*_1_,*𝒜*_2_) at time *τ* generations ago is bivariate normally distributed with covariance *Tσ*^2^, variance *τσ*^2^ and mean (*ℓ, ℓ*), the probability density of which we denote *f*_*t*_(*x*_1_, *x*_2_).

The probability that both lineages are to the right of zero *τ* generations ago, and hence are both of ancestry *B*, is therefore given by:

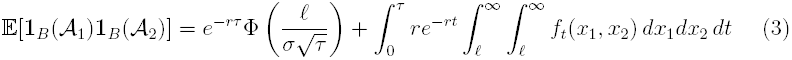

The first term of Eq. 3 corresponds to the probability that there is no recombination multiplied by the probability that the path of our single ancestral lineage is on the right side of the barrier when the barrier was removed. The second term integrates the probability that two lineages that recombined *t* generations ago are both to the right of of the barrier, i.e. the bivariate normal density integrated over the quadrant *x*_1_ > 0 and *x*_2_ > 0, over all possible times of first recombination. Rescaling *t* so that *u* = *t/τ*, equations 2 and 3 come together to give:

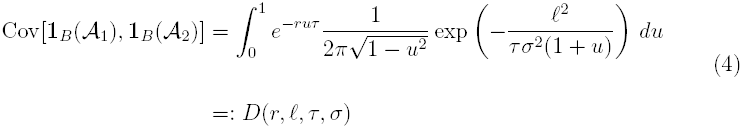

To obtain this expression, we integrate by parts, make use of the identity in Eq. A3, and rescale (0,*τ*) onto (0,1) (see Appendix A for more detail). We denote this covariance as a function *D*(*r, ℓ, τ, σ*), which expresses the expected covariance in ancestries of two loci in a randomly sampled individual from a given geographic location (*ℓ*) as a function of recombination fraction (*r*) between the loci, time since admixture (*τ*) and rate of dispersal (*σ*). In Appendix B we also develop analogous results for arbitrary migration schemes in discretized space, for both continuous and discrete time.

### 2.4 Admixture block lengths

An extension to the above approach for describing admixture-LD between two loci is to consider how ancestry along the chromosome is partitioned into unbroken genomictracts of ancestry drawn from one parental population. This is a natural way to think about coancestry in admixed populations, and the genome-wide distribution of ancestry tract length can contain information about admixture.

We again examine a chromosome drawn at random at geographic position *ℓ*, this time considering the probability that between physical positions *P* and *Q*, separated by genetic distance *d*, the chromosome is only of ancestry *B*. As above, we assume that after linkage is broken by recombination, the products of recombination move independently with respect to each other. This again assumes that *τ* is small relative to the timescale of coalescence. Further, it ignores the correlation structure imposed by the pedigree (Liang and Nielsen 2014; Wakeley *et al*. 2012), the impact of which we return to in the discussion.

We note that our measure of recombination rate *d* will differ from the earlier definition of recombination fraction as we will be tracking all recombination events between *P* and *Q*. We now assume that recombination events occur as a Poisson process with rate *d*, which reflects genetic distance on the genetic map between our two endpoint loci, and assume no chromatid interference.

If there have been *K* recombination events that occurred along the tract of chromosome over the last *τ* generations, then this region has *K* + 1 genetic ancestors from time *τ* that have spatial locations **X** = (*X*_1_, …, *X*_*K*+1_). As we neglect coalescence, we assume these ancestors are distinct. The segment contains only ancestry from population *B* if all *X*_*i*_ > 0 (i.e. all *K* + 1 ancestors are to the right of 0 at time *τ*, see Fig. 1 for an example of K=2). We denote the probability of our segment containing only ancestry from population *B* as:

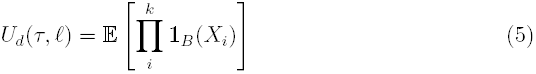

This is the expected value averaging over both the number and timing of recombination events, and the locations of the ancestral lineages at time *τ* ago (denoted **X**). We now outline one approach to obtain an expression for *U*_*d*_(*τ, ℓ*), and give a complementary approach in Appendix D.

#### 2.4.1 Obtaining block length distributions by summing over the number of recombination events

Since we assume no coalescence, the branching order of the ancestral lineages via recombination specifies a labeled tree structure, **S**, with *K* + 1 tips and a vector of splitting times **T** = (*T*_1_, …, *T*_*K*_) (where these times satisfy the constraints imposed by the tree topology). Since, looking backwards in time, each lineage moves as an independent Brownian motion once it has split from the others, the (*K* + 1)-lengthvector **X** of geographic positions at time *τ* is distributed as a (*K* + 1)-dimensional multivariate normal with mean (*ℓ*, … *, ℓ*) and variance-covariance matrix Σ. The entries of Σ reflect the shared path of tips *i* and *j*, so that 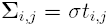, where *t*_*i,j*_ is the time of the recombination that separates tip *i* from tip *j*, and the diagonal entries 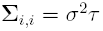. Conditioning on *K* = *k* recombinations and the matrix Σ, the probability that all *k* + 1 tips are of ancestry *B* is given by the integral of the *k* + 1-dimensional normal density over the space for which all *X*_*i*_ > 0:

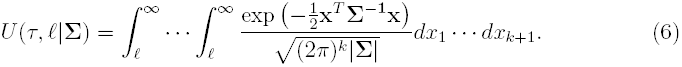

The integrand is the density for the multivariate normal which is determined by the timing and ordering along the chromosome of recombination events.

This needs to be averaged over possible trees; to do this, we sum over possible tree topologies, and for each tree topology integrate over possible split times (*T*_*i*_ ∈ [0, *τ*]). For a given tree topology *𝒯*, the term we need is the following (also rescaling spatial and temporal variables so that 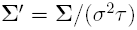 and 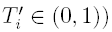:

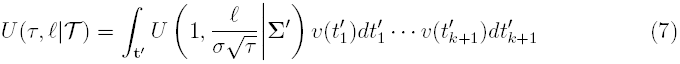

The set of possible times, **t**^′^, over which we integrate depends on the tree topology, and correspondingly, each topology has a weight, or probability conditioning on *k* recombinations that is given by 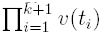 (See Appendix C for a further description of **t**′ and *v*(**t**′).)

Finally, we sum across *k* and 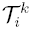 in the set 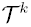 of all topologies given *k* recombination events.

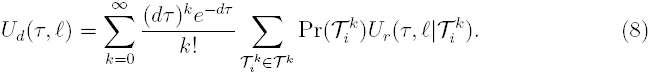

Where 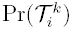 is the probability of the *i^th^* unlabeled topology given that there are *k* + 1 tips (we describe the calculation of 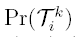 in the Appendix C.) We note that Eq. 8 is a Wild sum expansion for *U*_*d*_(*τ, ℓ*) (Etheridge 2000). We outline an approach using differential equations to obtain an equivalent expression in the Appendix D.

In practice, we approximate this sum by conditioning on *k*^*^ or fewer recombination events in *τ* generations:

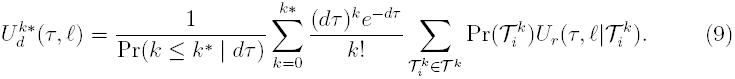

Fig. 2 shows the convergence as *k*^*^ is increased, to the distribution of tract lengths obtained by simulating *under the model* (see below for description for simulations under the model). Summing over the large number of topologies for large *k*^*^ is computationally expensive, but terms in the sum can be reused over some parameter values.

**Figure 2:**
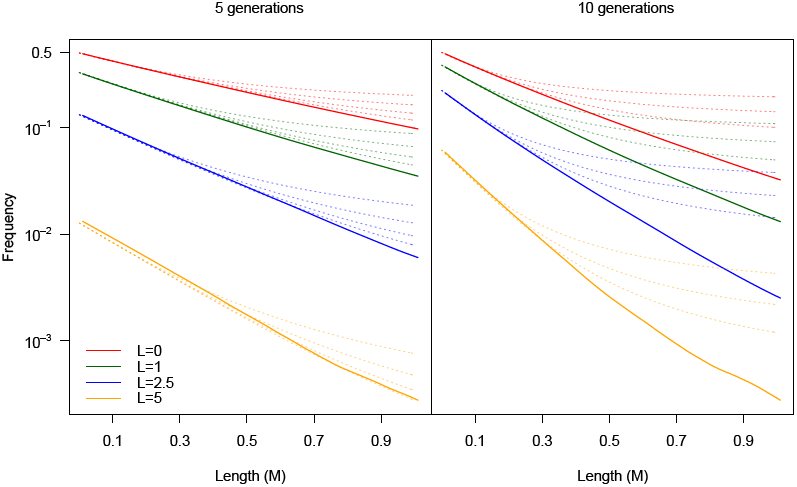
Distribution of tract lengths, expressed as the frequency of tracts that are at least a given length (i.e. 1-cumulative distribution of tract lengths). The following shows the distribution for populations *L* units away from the center of a contact zone. The solid lines represent the output of a simulated contact zone with no drift. For the 5-generation contact zone the four dotted lines per geographic position represent the predicted distribution under approximations conditioning on at most 3,4, 5 or 6 recombination events. For the 10-generation contact zone, the three dotted lines represent approximations conditioning on at most 3,4 or 5 recombination events.

### 2.5 Simulations

We developed two classes of simulations to (*1*) evaluate the accuracy of our analytic results, and (*2*) to explore the consequences of realistic violations of our model that likely occur under the specified biological process. For the first class of simulations, **simulations under the model**, we consider chromosomes moving in continuous space and time, with recombination modeled as a Poisson process through continuous time and independent movement of all products of recombination. This is an explicit simulation of the model described above. We simulated 10000 chromosomes *under the model*.

In the second class of simulations, **simulations under the process**, we follow a finite number of chromosomes migrating across discrete demes with non-overlapping generations forward in time. In these simulations we maintain the complete recombination history of a chromosome. As these features allow genetic drift, enforce a pedigree structure onto local ancestry, and occur in discrete time and space, our simulations under the process present a biologically realistic challenge to many of our major modeling assumptions. We consider 200,000 diploids (400,000 chromosomes) evenly spread across 20 demes. Demes are connected through nearest-neighbor migration with a per-generation, per individual probability *m* of migration (this migration rate is reduced to *m/*2 on demes at the edges of one-dimensional space). We sample the number of recombination events from a Poisson distribution with mean of one, corresponding to a 1 Morgan chromosome, and recombination events are uniformly placed along a chromosome (i.e. no recombinational interference). Every generation, migration, random mating, and recombination take place, and we follow the positions of all tracts of ancestry. After *τ* generations, we can sample chromosomes where the ancestor from the initial population of each locus is known. We then assign ancestry along each individual’s chromosome based on whether ancestors originated in population 1–10 (ancestry B) or in populations 11–20 (ancestry A).

### 2.6 Inference of parameters in human admixture data

While the distribution of continuous-ancestry tracts necessarily contains more information than LD alone, there are limits to the precision of the measurement of tract length over short recombination distances (which would reflect old events). This, combined with the relative ease of obtaining LD measurements from genomic data, motivates our use of LD in our analysis of human admixture contact zones. A variety of methods, including ALDER (Loh *et al*. 2013) and Globetrotter (Hellenthal *et al*. 2014) estimate some measure of ancestry-LD. We use the weighted LD curves generated by ALDER, which estimates a quantity analogous to the covariance in ancestry by computing the statistic:

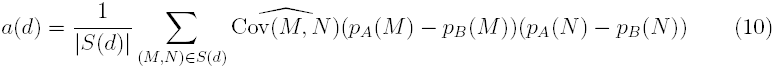

for a set of pairs of autosomal loci, *S*(*d*), that are a genetic distance *d* apart.

Here, (*M*, *𝒩*) is a locus pair, *p*_*𝒜*_(.) and *p*_*B*_(.) are sample allele frequencies in the parental populations *A* and *B*, and 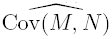 is the sample covariance between alleles at the two loci within the target population. If *r* is large enough that background LD in the ancestral populations can be ignored, and that the allele frequencies in the parental populations are known, then 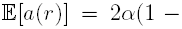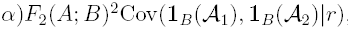, where 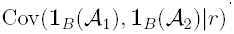 is the expected covariance in ancestry between pairs of loci a recombination fraction *r* apart *α*, is the ancestry proportion of population *𝒜* in the admixed population, and the constant *F*_2_(*A*; *B*)^2^ measures differentiation in allele frequency between the two parental populations. Often, the designated parental populations for analysis are proxies for the true parental populations, in which case *F*_2_(*A*; *B*)^2^ is a measure of the differentiation between the true parental populations that is shared by the proxy populations.

#### Admixture at a single-time point

Under a basic model of admixture, decay in ancestry-LD can be described by the parameters *F*, *t* and *G* in the exponential model

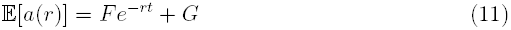

corresponding to a single pulse of admixture *t* generations ago. The term, *G*, represents admixture LD between unlinked markers, possibly due to substructure in the sampled individuals with respect to their ancestry proportions. The value *F* + *G*/2 corresponds to 2α(1 − α)*F*_2_(*A*; *B*)^2^ (Loh *et al*. 2013), where *α* is the admixture proportion, and therefore is a compound parameter reflecting both admixture proportion and differentiation between parental populations.

#### Fitting to a geographic contact zone

We take a set of admixed samples drawn from *n* populations, who fall at positions *ℓ*_1_, …, *ℓ*_*n*_ along a linear geographic transect. The geographic location of the center of the zone along this transection is *C*, such that sample 1 is a distance *ℓ*_1_ − *C* from the zone. We specify a pair of proxy parental populations *A* and *B*, to represent the end points of the contact zone. Using ALDER we generate the statistic *a*_*j*_(*r*_*i*_) for the *j^th^* population sample for each genetic distance bin (*i*), giving us a set, a, of weighted-LD decay curves (as defined in Eq 10). We use the minimum inter-SNP distance determined by ALDER based on LD in the parental populations.

To assess the uncertainty in a, we estimate the variance in ALDER’s statistics using the jackknife (which is an output of ALDER). For each of the *c* = 22 iterations, one chromosome is removed before recalculating a for the remaining 21 chromosomes. We use this to calculate the variance 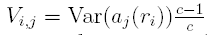. We then conduct a least squares fit of the ALDER output to our prediction given by Eq. (4) for values of *τ*, *σ*, *F* (corresponding to *F* in Eq. 11 and *C*. We fit all *n* populations simultaneously), calculating:

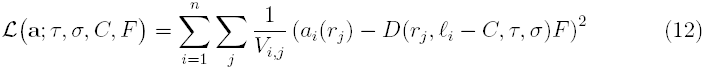

Our choice of 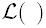 would be the negative log-likelihood of our parameters if our *a*_*j*_(*r*_*i*_) were normally distributed, a reasonable approximation given the large number of pairs of markers contributing to each value of *a*_*i*_(*r*_*i*_). We refer to 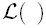 as the log-likelihood, and because we are mainly interested in *τ* and *σ* we generate profile surfaces of *τ* × *σ*. Specifically, we set a value for *L* based on a fit of Eq. 1 to ancestry proportion, generate a likelihood surface over a grid of *τ* × *σ* × *F* and for each combination of *τ* and *σ* we defined the profile log-likelihood as the maximum log-likelihood across all of our coresponding *F* grid-points.

We note that, although Eq. 11 includes an affine term to account for LD that could be generated by an unspecified model of population substructure, our model does not. This is because a source of long-range LD is incorporated into our model via gene flow from neighboring populations with different admixture proportions.

## 3 Results

### 3.1 Simulation results and comparison to exponential model

Figure S1 shows the decay in LD at various points in time and space, and shows the exact correspondence between the analytic expression of Eq. 4 and the output of *simulations under the model*. To evaluate the consequences of of fitting a single pulse model to data generated by our spatial model of continuous admixture, we fit the exponential decay of Eq. 11 to a set of simulated populations from a 50-generation old contact zone. The comparison, shown in Fig. S2, of best fit parameters indicates that the simple exponential tends to underestimate the age of the admixed populations by as much as a factor of 2, presumably because of the continuous introduction of migrants bearing long ancestral haplotypes. In other words, the poor fit of the single pulse model to these LD decay curves, especially close to the center of the contact zone, is due to the heterogeneous mixture of recombination times. Consistent with this interpretation, the effect diminishes in populations far from the center of the zone, as the difference in ancestry composition between neighboring populations decreases as the distance to the center increases.

To demonstrate our inference method as described above, we fit our model (Eq. 4) to the curves generated *under the process*. Because we simulated single chromosomes, we could not use the jackknife estimator of variance, and therefore modified Eq. 12 by removing the denominator. We removed populations with no detectable admixture from the fit, limiting our analysis to populations close to the center of the contact zone. The profile likelihoods of these surfaces are shown in Fig. 3). The inferred *τ* and *σ* are (2, 0.17), (38, 0.12) and (93, 0.11) for zones simulated under *τ* = 5*, τ* = 50 and *τ* = 100 respectively, under *σ* = 0.1

**Figure 3:**
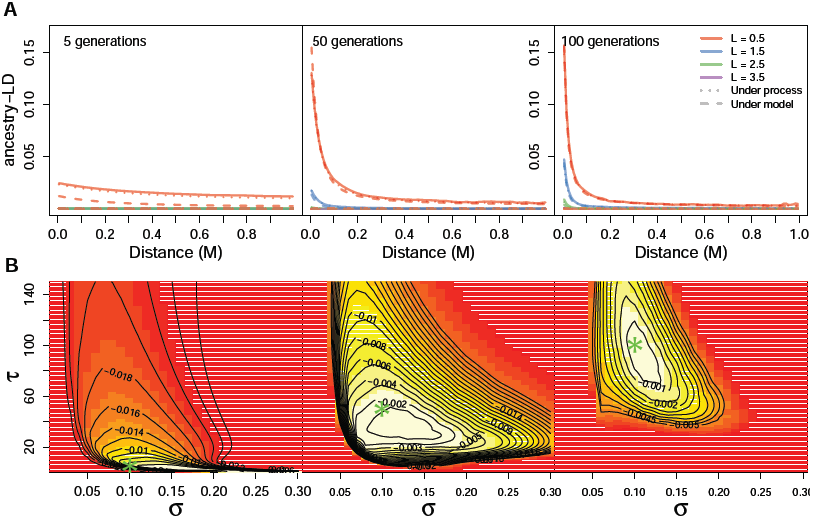
Analysis of simulations *under the process* run with parameters *τ* = 5, *τ* = 50 or *τ* = 100 and *m* = 0.01 under nearest-neighbor migration, corresponding to *σ*^2^ = 0.01 in the continuous model. **A:** Output of simulations (solid lines), compared to the continuous time and space model of Eq. 4 (dashed lines) and a discrete time and space expression from Eq. A7 (dotted lines). **B:** Profile likelihood surfaces describing the fit of our continuous model to simulations *under the process*. Green asterisks indicate simulated values.

Compared to the true values we use to *simulate under the process* our inference method tends to slightly underestimate the age of the contact zone. We expect that this is in part due to the discrete nature of the simulation. These estimates are closer to the true simulated ages than those obtained by fitting an exponential (Eq. 11) to each population, which return values of 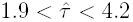 for *τ* = 5, 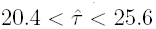 for *τ* = 50 and 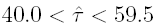 for *τ* = 100 compared to our values of 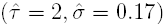 for (*τ* = 5, *σ* = 0.1), 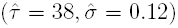 for (*τ* = 50, *σ* = 0.1) and 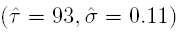 for (*τ* = 100, *σ* = 0.1).

### 3.2 Application to human datasets

We applied our model to three independent sets of populations that potentially represent admixture in a spatial context: Populations along the Indonesian archipelago, populations in Central Asia and populations in India (Table S1). Genetic distances between SNPs were inferred using sex-averaged recombination rates from deCODE (Kong *et al*. 2010).

#### 3.2.1 Indonesian archipelago

Populations along the Indonesian archipelago show a longitudinal cline of admixture between East Asian and Papuan autosomal ancestry (Xu *et al*. 2012; Lipson *et al*. 2014; The Hugo Pan-Asian SNP Consortium 2009). The decrease in proportion of Asian ancestry with longitude has been interpreted as evidence of the Austronesian expansion from the West through Indonesia. Xu *et al*. (2012) fit simple admixture models independently to each of the populations to infer admixture times of 120–200 generations, such that populations with higher Papuan ancestry have more recent admixture times. A more recent analysis using ALDER estimated single admixture dates for populations in the region in the range of 30–60 generations, suggesting that this in part is the result of subsequent waves of gene flow from populations with varying levels of Asian ancestry (Lipson *et al*. 2014).

We obtained the genotypes for seven population samples in Indonesia (shown in Table S1) from The Hugo Pan-Asian SNP Consortium (2009) and a Papuan population from the HGDP dataset (Li *et al*. 2008). We first ran STRUCTURE (Pritchard *et al*. 2000) with *k* = 2 on these nine samples. The admixture proportions obtained from STRUCTURE confirm the east to west cline (shown in Fig. 4). We then ran a least squares fit for Eq. 1 on these admixture proportions, which estimated the cline center at *X* = 124°9′E and *σ*^2^*τ* = 50.9. Based on ancestry proportions, we chose the Mentawai population and the Papua New Guinean population (with ~55k shared SNPs) as proxy source populations to generate ALDER curves. Simultaneously fitting our model to the six admixed populations, we generated the profile-log likelihood surface shown in Fig. 4. The maximum likelihood parameters best fitting these curves were an approximate contact time of ~200 generations or 5800 years ago (given a generation time of 29 years, Fenner 2005), *σ* = 0.63 and *F* = 0.0045. The fit to LD decay curves under these estimates is shown in Fig. 4 and Supplementary Fig. S3.

**Figure 4:**
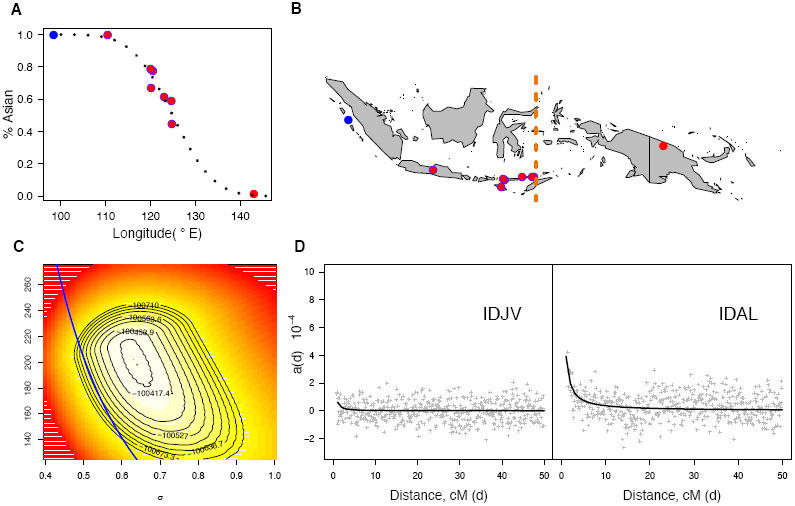
**A:** Longitudinal cline in Asian ancestry. Black dotted line shows best fit to Eq. 1. **B:** Sampling locations of Indonesian populations. Blue dot denotes the representative Asian ancestral population and red dot the representative Papuan population. Vertical yellow line shows location of the inferred cline center. **C:** Profile likelihood surface for *τ* and *σ* under Eq. 12 for all admixed Indonesian populations. The blue line represents the curve 50.9 = *σ*^2^*τ*, corresponding to the value of this compound parameter that is obtained by fitting to admixture proportions alone as shown in Fig. 4A. **D:** Weighted-LD curves for two populations of different distances away from the center of the cline. Grey points represents estimates of LD generated by ALDER, and black curves are expected LD under the estimated parameters.

We also explored the fit to LD decay curves of the single pulse model, fitting Eq. 11 by least squares (weighted by jack knife variance as in Eq. 12). Unsurprisingly, the fit of our model is not as good as a model in which all admixed populations are considered as having a single admixture time but allowed different values of *F* (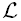 = 100370.1 compared to 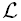 = 94147.7) since independently fitting the y-intercept to each population allows for many more parameters while these intercepts in our model are constrained by geographic distances between the populations. The fits to each population are presented in Table S2 and are in good accordance with those found by Lipson *et al*. (2014) using similar methods. With this approach, the mean timing among the admixed populations is 60.8 generations (we ignore the Javanese population which has little admixture and an estimated admixture time of 665 generations which as this is far older than all the other populations.)

Additionally, we considered fitting all populations simultaneously for a single time under the exponential model (Eq. 11), allowing each population to choose their own *F* parameter to account for differences in admixture proportions. Under this model we obtain an estimated age of τ ≈ 63 generations with minimum least squares of 98706. Again, this better fit is not surprising given that we are allowing each population to fit its own intercept.

Linguistic evidence suggests that the Austronesian expansion through Indonesia dates to ~ 2000 BCE (Gray *et al*. 2009). As noted by Lipson *et al*. (2014) these single pulse dates (Table S2) are too recent to reflect this, consistent with our earlier observation (and that of other authors) that admixture times may be underestimated by a simple exponential model if admixture has been ongoing. Our estimate of timing based on fitting a geographic contact zone (~200 generations) is much older than dates estimated by single pulse models, but is also considerably older than the Austronesian expansion. Considering that it is constrained by having to fit all populations simultaneously, our model provides a good fit. One possible explanation for our overestimate of admixture time is the assumption of a continuous rate of diffusion after initial contact. Despite this, our model may be a more realistic depiction of ongoing gene flow than a single pulse model and demonstrates that, in instances such as this where there is a gradient of admixture, incorporating a spatial model of admixture can provide additional insights into the history of these populations.

#### 3.2.2 India

Population structure in India is complex and multilayered. While the precise history of human movement in this region is unclear, work by Moorjani *et al*. (2013) and Reich *et al*. (2009) suggests that many modern Indian populations are descendants of an admixture event between differentiated Ancestral North Indian (ANI) and Ancestral South Indian (ASI) populations, with a cline in the extent of ANI ancestry across the subcontinent ((Moorjani *et al*. 2013), Fig. 5). While it is difficult to identify modern proxies of the parental populations, the ANI population appears to be most closely related to Western Eurasian populations (such as Georgia) and the Onge population of the Andaman Islands seem to draw much of their ancestry from the ASI population. Moorjani *et al*. (2013) broadly grouped their samples into Indo-European or Dravidian samples, and under this classification, found that the decay in ancestry-LD in their samples were consistent with two historical admixture events, one approximately 108 generations ago giving rise to the Dravidian populations, and a second wave of admixture from the north taking place 36 generations later that contributed to the ancestry of Indo-European populations.

**Figure 5:**
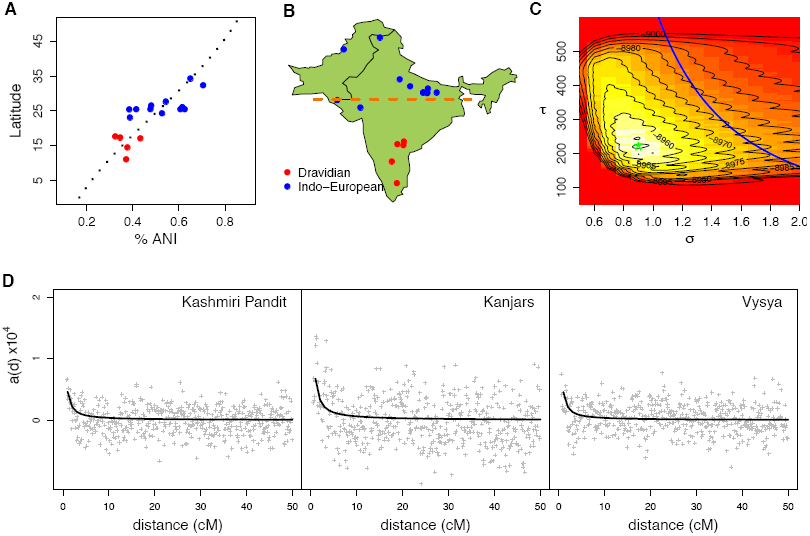
**A:** Latitudinal cline in ANI ancestry. **B:** Locations of Indian populations used in the analysis. Yellow line indicates location of inferred cline center. **C:** Profile likelihood surface for *τ* and *σ* under E1. 12. Blue line represents the relationship 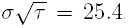, as obtained from the cline in ancestry proportion. Asterisk denotes values providing best fit. **D:** Weighted LD curves as estimated by ALDER, for a northwest (Kashmiri Pandit), southern (Vysya) and northeast (Kanjars) population. Grey points are estimates generated by ALDER, and black curves are expected LD under the estimated parameters.

We obtained the genomic data used in Moorjani *et al*. (2013), Reich *et al*. (2009), Metspalu and Romero (2011) and Li *et al*. (2008), yielding approximately 83, 000 shared SNPs, and focus on the populations represented in Table 1 of Moorjani *et al*. (2013) (See our Table S1). Following Moorjani *et al*. (2013), we ran the *F*_4_ ratio tool in the ADMIXTOOLS package (Patterson *et al*. 2012) on Georgian, Basque, Yoruba, Onge and the focal Indian population to estimate ANI ancestry proportions in these populations (Fig. 5). We fit a latitudinal cline to these ancestry proportions (Eq. 1) returning a cline center at 24°4′*N* and 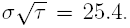. Because the gradient of ancestry could run along any geographic axis, we also tried to fit ancestry proportion clines to various transects using linear combinations of latitude and longitude. Since these did not produce substantially better fits than latitude alone, we chose to use latitude as our geographic axis (results not shown).

We then generated co-ancestry decay curves in ALDER for each of these samples, using weightings from Basque and Onge parental populations as proxies for the ANI and ASI populations, see Moorjani *et al*. (2013). We consider three possible contact zone scenarios: One in which all population samples form a contact zone and, based on the earlier studies, one that comprises only of the Indo-European and one that comprises only of the Dravidian populations. We initially attempted to fit the *τ*, *σ* and *F* parameters in Eq. 12 simultaneously, but faced some difficulty as there appears to be limited information about *F*. This results in wide range of values fitting the data equally well, but give rise to very different surfaces for *σ* and *τ*. We attributed this to a deficit of information in the curves, leading to non-identifiability, due to relative low levels of differentiation and relatively rapid decay of ancestry-LD. The difficulty in estimating the intercept of admixture-LD curves had been noted before (Loh *et al*. 2013), and can reflect the fact that very close pairs of markers are discarded to remove the effects of LD in the ancestral populations. This results in the fitted curve being relatively unconstrained near *r* = 0. To remedy this, we estimated *F* using an approach similar to that taken by Moorjani *et al*. (2013). Using MIXMAPPER (Lipson *et al*. 2013), we estimated the value of *F* as *F*_2_(*ANI*; *ASI*)^2^ using the Onge and Basque populations as present day proxies, and fit values of *σ* and *τ* under the range of *F*_2_ values computed by MIXMAPPER ((0.015, 0.042)). We also use the value estimated above as the cline center for all three fits. We first fit our LD curves to all populations, under a model in which all Indo-European and Dravidian populations are the outcome of a single admixture contact zone. The best fit was approximately 220 generations since contact (Fig. 5). Fits to the subset of populations classified as Indo-European yielded a contact zone age of approximately 200 generations (Fig. S5). Finally, we fit the subset of Dravidian populations (Fig. S5), which found a best fit of 460 generations on a relatively flat surface. This is likely because there is very little information in the decay of LD in this subset given there are so few Dravidian populations, and that the LD curves are relatively flat.

Several aspects of the data indicate potential mis-estimation of dates. Some populations, presumably the oldest, have very little admixture-LD, which may prevent an accurate fit to the decay. Secondly, it is possible that the absence of ‘edge’ populations that are further away from the zone center makes it difficult to obtain a good fit, as we only have populations with intermediate levels of admixture where the decay of LD is not strongly related to the age of the zone. Substructure within populations, due to practices such as endogamy, may also influence ancestry-LD within a population and cause a deviation from expectations under a null model of randommating. We take these challenges, and the uncertainty in our results, as a reflection of the complicated demographic histories of these populations, and the fact that it is poorly described by the model which we are trying to fit. These challenges also likely apply to other analyses of these data, meaning that caution is warranted in judging the age of this zone.

#### 3.2.3 Central Asia

Populations in central Eurasia show varying levels of East Asian ancestry. In a global analysis, Hellenthal *et al*. (2014) identified a signal of admixture, using Mongolian and Iranians as proxy source samples, in Turkish, Uzbek, Hazara and Uygur samples. The proportion of Mongolian ancestry decreases with longitudinal distance from Mongolia, with the Turkish populations harboring the lowest proportion of Mongolian ancestry. The estimated admixture dates in these populations of 20-30 generations in the past found by Hellenthal *et al*. (2014) is consistent with the timing of the westward military movement of Mongolians during the 13th century.

We took the genomic data for the four admixed populations and the two proxy source populations from the dataset of Hellenthal et al (500k SNPs). A STRUCTURE analysis of these populations, with *k* = 2, is consistent with a gradient in Mongolian ancestry across Central Asia (Fig. 6). We used ALDER to generate weighted covariance curves, using the Mongolian and Iranian samples as the two proxy source populations. For the four admixed populations, the best fit (Eq. 18) under our simple contact zone model is approximately 49 generations, or 1421 years ago (29 years per generation), with *σ* = 3.7 (see Fig. 6 for the profile likelihood surface). This admixture date predates the Mongolian invasion of Central Asia that took place approximately 800 years ago. However, it is known that human movement in Central Asia was complex, and preceded the Mongolian invasions by centuries, and it is possible that our estimated date is capturing a signal of these earlier migrations. This is supported by recent analyses of Central Asian populations by (Yunusbayev *et al*. 2014).

**Figure 6:**
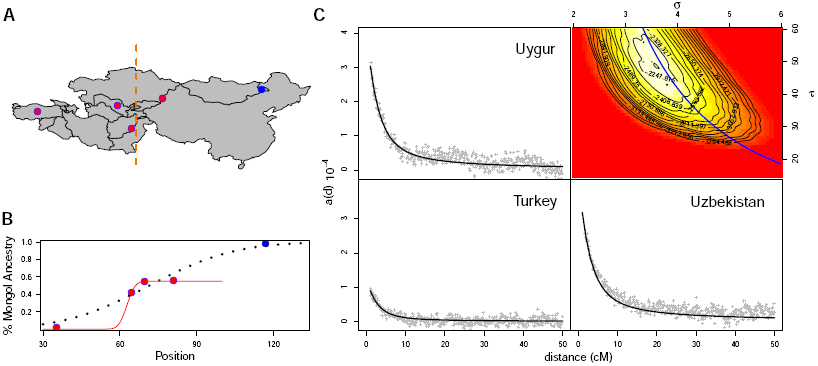
**A:** Geographic location of Mongol-Iranian admixed populations used in the analysis. **B:** Ancestry proportions, with best fit under basic Brownian model (dashed, thick line), and under pulse model (unbroken thin line) C: Best fit under our model to LD-decay curves (Hazara not shown), and profile likelihood surface to the set of all four populations (top right). Blue line indicates 4.2 = *σ*^2^*τ*, the compound parameter estimated by fitting to admixture proportions.

ALDER identified a large extent of long-range LD in the Hazaran population, possibly due to population substructure within this sample with respect to Mongolian ancestry. Because this could potentially influence our inference, we refit the LD curves to the set of admixed populations exculding the Hazara. This produced a best fit of 37 generations.

One consideration in our applications is our assumption that the populations spread back into contact and then simply passively diffused into each other. This is obviously likely a poor description of the movement of Mongolian genotypes across Asia during the 13th century invasions, which could result in a discrepancy between expected and predicted decay in ancestry-LD. We therefore proposed an alternate model that allows for an initial fast pulse of Mongolian migration into central Asia, followed by diffusion through local geographic dispersal (i.e. our Brownian motion). Explicitly, we construct a model which defines two additional parameters: *X*_1_, a point in space to the east of which some proportion, Ψ, of the population is replaced by Mongolian genotypes *τ* generations ago (see Appendix E for mathematical details). In specifying this model, we are trying to capture a scenario in which, at least initially, unadmixed Mongolian genotypes were making a rapid westward movement. However we acknowledge that this is at best a very crude approximation of a possible sequence of events.

While this alternate model provides a better fit to admixture proportions (Fig. 6 shows fit with Ψ = 0.55 and *X*_1_ = 62.7), given the few populations, this good fit may reflect over-parameterization of the model. Furthermore, a search for the best fit to the LD decay curves returned parameters that were effectively identical to the initial basic model proposed (Ψ *≈* 1, cline center around 71°*E*), indicating that this is not a likely alternative model (profile likelihood curves for each fitted parameter are shown in supplemental figure S8). Given the early estimated admixture date, it is possible that admixture across Central Asia is not a product of a single event as our models, and those of others (Hellenthal *et al*. 2014), assume, but rather a result of complex human migrations throughout time. Despite the limitations imposed on inference of parameters by the small number of populations, broad patterns of ancestry-LD across space are nevertheless somewhat consistent with our proposed model of ancestry-LD decay across space along an admixture gradient.

## Discussion

The generation and subsequent of decay of admixture-LD as an outcome of inter-breeding between differentiated populations provides a population genetic signature that is a valuable tool for understanding the nature and timing of admixture. Existing methods for modeling decay in admixture-LD consider the expected rate of decay in one population at a time, and often assume a simple one-time ‘pulse’ of admixture without subsequent gene flow from neighboring admixed populations. Here, we have described a neutral model under which individuals diffuse across space. Based on this model, we derive an analytic expression for the expected decay in ancestry-LD as a function of time since contact and a population’s position in space. We consider this an alternate model to one in which admixed populations are independently formed by a single-pulse event with potential subsequent gene flow from parental populations. In contrast to previous analyses of spatial admixture which treated populations as independent admixture events (e.g. Xu *et al*. 2012), we consider data from all sampled populations simultaneously to build a model that incorporates all available information and accounts for the movement of individuals between populations. Compared to the expression for ancestry-LD derived here, a simple exponential model tends to underestimate the time since admixture, as it does not account for the introduction of long ancestral haplotypes from neighboring populations.

### Additional sources of covariance

In developing tractable approximations to spatial admixture contact zone we have ignored genetic drift and the genealogical structure imposed by the pedigree.

Genetic drift is not problematic if population densities, and dispersal rates, are high enough that coalescence between geographical close lineages is unlikely over the time-scale *τ* (as is likely the case in our human applications). Otherwise, a theoretical approach incorporating coalescence will be needed (see Barton *et al*. 2013, for recent progress). However, in that case, background LD and admixture LD will be on comparable genomic scales, making the the job of separating the two much more challenging.

The other form of correlation structure that we have ignored is that imposed by the genealogy (Wakeley *et al*. 2012; Liang and Nielsen 2014). When there are multiple crossovers during meiosis within the stretch of chromosome we are considering, the recombinants trace their ancestry to one of the two parents one generation back in time. When considering the chromosome tracts between recombination events, odd numbered recombinant segments come from one parent (say the mother), and even number segments from the other parent (the father). Therefore, the recombinants are not independent of each other as one generation back as all odd (or all even) recombinants are found in one parent. This additional covariance from the pedigree structure does not impact our pairwise model of ancestry-LD if *r* is strictly defined as a recombination fraction, as an odd number of recombinations between our pair of loci means that the two alleles are present in different parents in the proceeding generation and there after follow independent trajectories back in time. Our block length calculations ignore this form of covariance, as we assume that fragments follow independent spatial paths backward in time after recombination events. This assumption will only be problematic for long regions (where more than one recombination can happen per generation) and for short time intervals (i.e. small *τ*). However, in such cases, ignoring genetic interference may present a greater source of error than the ignoring of this additional source of covariance.

### 3.3 Application of the model to human admixture data

To explore our model we used our approximate model to estimate contact times and dispersal variance from genomic data from admixed human populations. We present our fit to the output of weighted-LD from ALDER, but similar information about the extent of ancestry-LD can be obtained from alternative methods such as Chromopainter (Lawson *et al*. 2012).

Our spatial model provided a good fit to admixed populations along the Indonesian archipelago, consistent with a relatively straightforward history of admixture across space. Our estimated time of initial contact is somewhat consistent with the work of Xu *et al*. (2012), and is older than reported by Lipson *et al*. (2014). Our deeper admixture time estimate likely reflects the fact that inference under single-population admixture models will produce estimates of timing of initial admixture that is more recent than estimates under our contact zone model. Our estimate of *≈* 6000 years ago is older than estimates obtained from linguistic analysis (Gray *et al*. 2009). This could be in part due to the simplifying assumptions of our model, which requires dispersal to be constant in time and space. One could imagine, for example, that if there were pulses of human movement followed by a slowing down of dispersal this would impact our estimate.

Our spatial model provided a poor fit to the Indian and Central Asian populations. This is likely due, in part, to deviations from a simple model of instantaneous removal of a barrier to contact and continuous diffusion thereafter. In India, a complex population structure, caste system, and potentially two waves of contact may have all contributed to difficulties in finding parameters that fit under our model. In particular, the need to separately estimate the y-intercept meant that there was relatively little information in the decay curves about the timing and mode of admixture. This is especially problematic for older admixture such as this (particularly in the Dravidians), as there is relatively little admixture-LD over larger scales and consequently much of our information relies on LD over short genetic distances (*<* 1cM). Given this paucity of information, it is likely that many, and quite different, admixture models would fit these data nearly equally well. As such, our fit and estimate of timing, and indeed the estimates under alternate models, should be interpreted with caution.

The limited number of populations in Central Asia places a limit on the confidence for the fit to the data under any dispersal model. Furthermore, it is known that human movement in the region spans many centuries and is unlikely to be simple. While earlier attempts to date admixture in these populations estimate admixture times of *≈* 30 generations, corresponding to the Mongolian invasions (Hellenthal *et al*. 2014), our estimated time is much older, at *≈* 50 generations. It is unlikely that our demographic model is a good approximation to historical human movement in the area, and this is likely to have impacted our inference. However, it is possible that our estimate of earlier admixture is in part reflecting older human movements in the region, and this is in part supported by the findings of (Yunusbayev *et al*. 2014).

### 3.4 Extensions of the simple neutral model and other applications

The assumption of Brownian movement, and the ignoring of drift and pedigree structure have enabled the derivation of a relatively simple expression to describe ancestry-LD. The examples of human admixture zones provides above indicate, however, that alternative models may be need to describe patterns of LD, given different demo-graphic scenarios. We therefore consider the basic Brownian model to be a neutral framework and acknowledge that, while it may be a good approximation for some scenarios of admixture and secondary contact, in many cases individuals may not diffuse continuously in space and time. Because of the simplicity of our model, modifications can be made with relative ease to describe different geographic scenarios. For example, we were able to apply a model in which the movement of Mongolian genotypes began as a pulse of migrants, followed by diffusion. In a similar vein, one could modify movement to contain a Brownian drift parameter to account for directional migration, although this would require some consideration as to how the dispersal kernel of an admixed individual is determined. Discrete deme models could also be used (as we develop in Appendix B) to model complex histories of populations in geographic and temporal heterogeneity. However, in practice there is not enough information in admixture decay curves to infer detailed population histories with many parameters.

We have demonstrated that inference of admixture parameters can be greatly influenced by the choice of demographic model. We believe that this highlights the need for more admixture models to be developed to test with population genomic data, and for careful consideration of which model is appropriate for a given biological scenario. The model presented here makes some progress towards addressing the movement of admixed individuals, and presents a potential framework for future development of dispersal models. As a final point, we note that all (to our knowledge) admixture models to date, including ours, assume that populations undergo differentiation in relative isolation prior to secondary contact. Under this assumption, there is a strong appeal to fit pulse models (such as a wave of secondary contact) to human admixture data, with a goal to estimating the timing of a pulse, and relating it to particular historical events. It seems that perhaps a more appropriate null model in these scenarios would be one in which gene flow has been ongoing between populations, but at a rate slow enough to allow some differentiation to occur. Testing for patterns of LD under this isolation-by-distance model would be a first step towards understanding the demographic history of spatially distributed populations, and the development of such a null model seems an important step in creating future tools for population genomic inference.

In addition to admixture contact zones, LD has been used to characterize hybrid zones (Wang *et al*. 2011), and we see our framework as a potential null model for spatial models of secondary contact, whereby incipient species come back into contact. Although tension zones can maintain distinct species, reproductive isolation is often weak enough to allow diverged populations to exchange alleles. In such scenarios, patterns of diversity that depart from expected ancestry-LD could be used to detect potential targets of selection relevant to speciation or local adaptation. The expected population genomic signatures of such loci will depend on the nature of selection – for example, patterns of LD around a gene under differential selection may differ from patterns of selection against certain hybrid genotypes. It should be noted, however, that good estimates of decay in ancestry-LD require reliable genetic maps, as overestimates of genetic distance may give the appearance of a slower rate of decay by inflating LD and this may be a limiting factor in many systems.

The LD induced by the admixing of two differentiated populations is a powerful population genetic tool which, combined with genome-wide data, has enabled the use of decay in ancestry-LD to inform the timing of admixture events. Building on models that use this decay to infer admixture dates under scenarios with discretized migration events, we have developed a novel framework that accounts for continuous movements of haplotypes through time and space. We believe that this can serve as a good null model for understanding patterns of diversity in contact zones. Further-more, we see potential for this model to be further developed and tailored to fit a range of demographic scenarios, including those that incorporate selection.

## Acknowledgements

We thank P. Moorjani, G. Hellenthal and M. Metspalu for access to data, and Simon Aeschbacher, Alison Etheridge, Jeremy Berg, Gideon Bradburd and Ivan Juric for helpful conversations and comments. This work was supported by the NSF GRFP under Grant No. 1148897 and by grants from the National Science Foundation under Grant No. 1262645 to P. Ralph and G. Coop and the National Institute of General Medical Sciences of the National Institutes of Health under award numbers NIHRO1GM83098 and RO1GM107374 awarded to G. Coop.

## Appendix

## A Covariance in Ancestry

By integration by parts, equation (3) becomes:

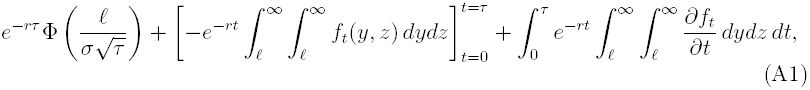

where *f*_*t*_(*y, z*) is the bivariate normal density for jointly distributed (*Y, Z*) with correlation *t*. The second term of (A1) is:

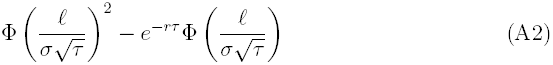

For the third term of (A1), we can utilize the useful identity that for a bivariate normal with variances 1 and correlation *t* (Pearson 1901):

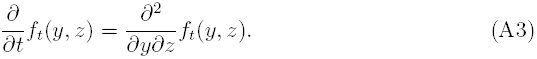

The last term of (A1) becomes:

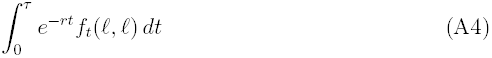

Combining Eq. 2 and A1, A3, A4 therefore leaves us with:

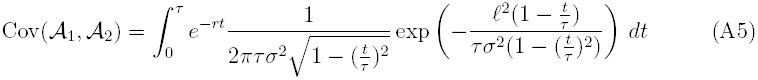

## B Island model

In a discretized time and space model, with *n* islands and per-generation migration rates defined by the *n* × *n* matrix *M*, the expected frequency of ancestry *B* alleles in population *X* is

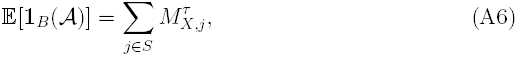

where *X* is the deme from which an individual is sampled, *τ* is the number generations since admixture began, *S* is the set of demes that are defined as being ancestry *B* at the time of contact, and 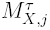 is element *i, j* of the *τ*^th^ matrix power of *M*. The covariance is derived by summing over possible recombination times and the location of the allele at the time of recombination (*𝒩* is the set of all locations.):

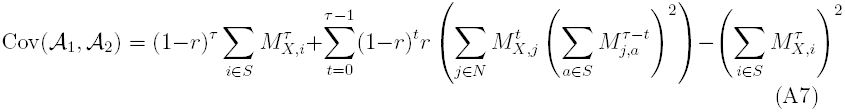

Note that *r* is the probability of any odd number of recombinations occurring, i.e. is the probability that a Poisson random variable with mean *d* is odd.

## C Unlabeled rooted topologies and their probabilities

To obtain the set 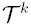 and the associated 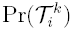, we use the following result from Cavalli-Sforza and Edwards (1967). Given *k* tips, the number of unlabeled topologies, *a*_*k*_ is given by the recursion:

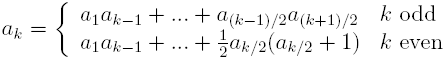

with initial conditions *a*_1_ = 1, *a*_2_ = 1

Intuitively, for the set of *a*_*k*_ topologies 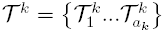, a topology 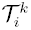 is generated by joining 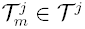 with 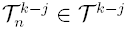 at the root (*m* and *n* are arbitrary). Because each subtree is independent, the probability of a topology given *k* recombinations can be calculated using a similar intuition. The probability, 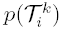, of topology 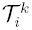 is the product of the probabilities of each subtree, relative to every combination that yields a tree of size *k*.

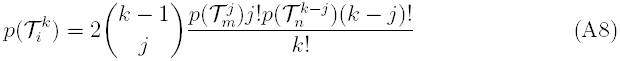

Where 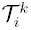 is the topology made by joining topologies 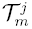 and 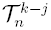 at the root.

The covariances for each topology representing *k* recombination events are dependent on the order statistics for *k* uniformly iid sampled recombination times. The **t**′ over which we integrate are conditional on these ordered recombination times. Specifically, if *t′*_*j*_ is the recombination time corresponding to node *j* on the tree, then *t′*_*j*_ becomes a lower bound for all subsequent recombination times associated with nodes that are descended from node *j*. Correspondingly, the factor 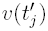 is a function of the recombination times

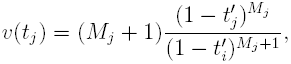

where node *i* is the parental node to *j* with corresponding time *t′*_*j*_, and *M*_*j*_ is the number of nodes descendent from node *j*. Here we have assumed recombination times are continuously distributed, and that double-recombination events do not occur (i.e. all nodes are unique with respect to timing.)

## D Obtaining block length distributions by a Branching Brownian Motion

An alternative approach the multiple-recombination scenario can be taken without conditioning on the number of recombination events. The process of recombination and dispersal described above is analogous to a Branching Brownian Motion (BBM), where recombination is represented by a splitting event. In standard BBM, lineages have a constant rate of splitting, but here the total length of the chromosome is constant, and so we have conservation of the total rate of splitting *d*. The rate of splitting on a lineage decreases with each recombination event, as both products of recombination are shorter (and therefore have a smaller probability of recombination).

Below, we derive an integro-differential equation satisified by *U*, similarly to the classic analysis of branching Brownian motion by McKean (1975). Starting in the present, we follow a single lineage backward in continuous time. The movement of this lineage is Brownian with variance *σ*^2^. We model recombination events between the two loci as a Poisson process with rate *d*. At the first recombination event, we generate a uniform random variable, *r*^1^ ∈ [0, *d*) to represent the genomic position of the recombination event. We then split the sequence into left and right fragments – [0, *r*^1^) and [*r*^1^, *d*), respectively. Following this, the two linages move independently backwards in time with respective recombination (splitting) rates of *r*^1^ and *d* − *r*^1^. This process is iterated over the time period *τ*.

We consider moving back a very short time interval Δ*t* from the present, and take the expectation over the random events that could have occurred in that time interval. (In other words, we are writing down the infinitesimal generator of this Markov process.)

With probability 1 − *d*Δ*t* + *O*(Δ*t*^2^) there is no recombination during the interval Δ*t* and conditioning on this, we have only to take the expectation over the small random change Δ*x* in spatial location during this time.

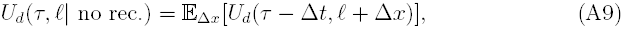

where 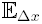 is the expectation over all changes in position *X*.

A recombination event occurs in the interval Δ*t* with probaility *d*Δ*t*. Conditioning on recombination occurring at time *t*_rec_ at position *ℓ* + Δ*x*′, producing two recombinants of length *d*_1_ and *d* − *r*_1_:

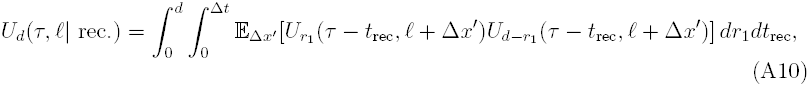

where 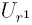 is the probability that all subsequent recombinants along the chromosomal fragment of length *r*^1^ are of ancestry type *B*.

As Δ*t* → 0, the Taylor expansion of (A9) and (A10) about *X* gives the expression:

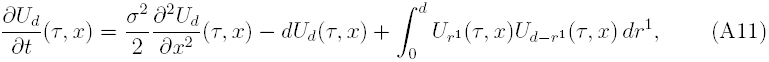

with boundary conditions *U*_*d*_(0, *x*) = 1 for *x* > 0 and *U*_*d*_(0, *x*) = 0 for *x* ≤ 0. This differential equation is solved by *U*_*d*_(*t, x*), defined in Eq. 5, and is the probability that at time *τ* in the past, the leftmost branch of this branching process initiated at position *x*_0_ is at a position *x* > 0. This differential equation is related to that presented by Baird *et al*. (2003) to describe the survival of genomic blocks within a panmictic population (but the latter does not have a spatial diffusion term). The equation is similar to the Fisher-KPP equation, with differences arising from the non-constant splitting rate. The first term of Eq. A11 reflects the spatial diffusion of lineages, the second term reflects the loss of blocks of length *d* to recombination. The final term reflects fact that the two recombinant lineages (of size *d − r*_1_ and *r*_1_) independently have to be of type *B*, and the dependence of this probability on the physical location of the recombination event, which is integrated over.

## E Invasion pulse

Suppose an invasive population displaces a subset Ψ of a resident population at *τ* generations in the past such that the frequency of ancestry *B* at time *τ* is 0 for −∞ < *x* < *X*_1_ and Ψ for *x* > *X*_1_. The ancestry LD at position *X* in this situation is:

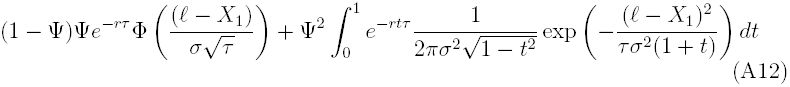

As after *τ* generations the probability of ancestry *B* is the probability of both of our lineages being in (*X*_1_, ∞) multiplied by Ψ

